# A genome wide search for non-additive allele effects identifies *PSKH2* as involved in the variability of Factor V activity

**DOI:** 10.1101/2024.02.08.579474

**Authors:** Blandine Gendre, Angel Martinez-Perez, Marcus E Kleber, Astrid van Hylckama Vlieg, Anne Boland, Robert Olaso, Marine Germain, Gaëlle Munsch, Angela Patricia Moissl, Pierre Suchon, Juan Carlos Souto, José Manuel Soria, CHARGE Hemostasis working group, Jean-François Deleuze, Winfried März, Frits R Rosendaal, Maria Sabater-Lleal, Pierre-Emmanuel Morange, David-Alexandre Trégouët

## Abstract

**Background:** Factor V (FV) is a key molecular player in the coagulation cascade. FV plasma levels have been associated with several human diseases, including thrombosis, bleeding and diabetic complications. So far, two genes have been robustly found through genome wide association analyses to contribute in the inter-individual variability of plasma FV levels: structural F5 gene and PLXDC2.

**Methods:** We used the underestimated Brown-Forsythe methodology implemented in the Quicktest software to search for non-additive genetic effects that could contribute to the inter-individual variability of FV plasma activity. QUICKTEST was applied to 4 independent GWAS studies (LURIC, MARTHA, MEGA and RETROVE) totaling 4,505 participants of European ancestry with measured FV plasma levels. Results obtained in the 4 cohorts were meta-analyzed using a fixed-effect model. Additional analyses involved exploring haplotype and gene×gene interactions in downstream investigations.

**Results:** We observed a genome-wide significant signal at *PSKH2* locus, on chr8q21.3 with lead variant rs75463553 with no evidence for heterogeneity across cohorts (p = 0.518). Although rs75463553 did not show association with mean FV levels (p = 0.49), it demonstrated a robust significant (p = 8.4 10^-9^) association with the variance of FV plasma levels. Further analyses confirmed the reported association of *PSKH2* with neutrophil biology and revealed that rs75463553 likely interact with two loci, *GRIN2A* and *POM121L12*, known for their involvement in smoking biology.

**Conclusions:** This comprehensive approach identifies the role of *PSKH2* as a novel molecular player in the genetic regulation of FV, shedding light on the contribution of neutrophils to FV biology.

## Introduction

Factor V (FV) is a central protein of the coagulation cascade. By acting as a co-factor for activated Factor X, FV facilitates the conversion of prothrombin to thrombin [1], the latter converting then fibrinogen into fibrin, the main component of blood clots and which also activates platelets. Mainly expressed in the liver, FV can also be stored and released by platelets [2] which provide a surface for the coagulation reactions to occur and which can contribute to amplify the coagulation process. There is natural variability in FV levels and increased/decreased FV levels have been observed in several conditions including mainly bleeding [3–6] and thrombotic disorders [7,8] but also in infections [9], inflammation [10], pregnancy [11,12], hormone contraceptives usage [13], and impaired liver dysfunction [14].

Understanding the exact sources of variability of FV levels is crucial for a better identification of individuals at higher risk of clotting disorders and for better targeting appropriate preventive and therapeutic strategies. Age, sex, smoking [15], obesity [16] and to lesser extent medication use [17], are the main environmental variables known to influence FV plasma levels. Genetic factors have also been demonstrated to contribute to the inter-individual FV variability including single nucleotide polymorphisms (SNPs) at *F5* and *PLXDC2* loci [18]. The implication of *F5* SNPs in the regulation of FV plasma levels dates back to the end of the 90s [19] where the HR2 haplotype tagged by rs6027 was identified. More recently, the *F5* rs4524 was also shown to influence plasma FV levels independently of the rs6027 [18]. The first genome wide association study (GWAS) on FV levels, based on ∼1700 individuals identified the *PLXDC2* locus as a second genetic player in FV regulation [18]. Altogether, these three loci explain less than 15% of the variability in plasma FV levels, suggesting that additional molecular determinants could be involved in its regulation. With the aim of characterizing novel genomic regulators of FV plasma levels, we here deployed a large scale agnostic genome wide search for non-additive genetic effects associated with FV plasma levels using the Brown-Forsythe (BF) methodology implemented in the Quicktest software [20]. While initially developed for detecting parental-of-origin effects (POE), this methodology can also detect loci prone to gene x gene or gene x environment interactions, making it a valuable tool to complement standard genome wide association analysis. POE is a specific kind of genomic imprinting [21,22] and several studies suggested that such epigenetic mechanisms could impact key genes of the coagulation cascade [23,24], including *PLXDC2* [25]. In this work, the BF methodology was applied to genome wide genotype data available in 4 study populations totaling 4,505 individuals with measured FV plasma levels.

## Materials & Methods

This work builds on four independent study populations of unrelated individuals, all of European ancestry, that are part of the *Cohorts for Heart and Aging Research in Genomic Epidemiology* (CHARGE) Consortium [26] : the *LUdwigshafen RIsk and Cardiovascular Health* (LURIC) study [27] composed of 1,833 individuals, the *MARseille THrombosis Association* (MARTHA) study [28] composed of 1,011 patients with venous thrombosis (VT), the *Multiple Environmental and Genetic Assessment* (MEGA) study [29] of 865 VT cases and the *Riesgo de Enfermedad TROmboembolica VEnosa* (RETROVE) [30] composed of a sample of 398 VT cases and of 398 controls. All participants were phenotyped for plasma FV activity and genotyped for genome wide polymorphisms using high-throughput DNA arrays. Genotype data were further imputed using different reference panels. Detailed descriptions of the phenotype and genotype measurements are given in Supplementary Table S1 together with additional details on imputation and genotype quality controls.

All participating studies were approved by the respective institutional Ethics Committees. Written informed consent was obtained from all participants to be included in such genetic investigations.

### Statistical Analysis

#### The Brown-Forsythe methodology implemented in the Quicktest program

Standard GWAS are generally performed for detecting SNPs having additive allele effects on a trait of interest. The statistical modeling can thus be expressed, in case of a quantitative trait Y, as

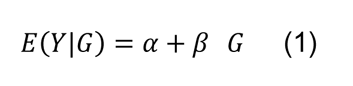

with G={0,1,2} according to the number of tested alleles carried by an individual. This model assumes the absence of POE for the tested allele while POE would imply that the effect of the tested allele would depend on whether it has been inherited from the father or from the mother. In that case, a POE model could be written as:

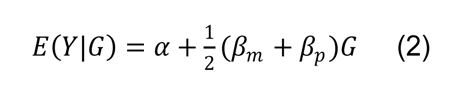

where β_m_ and β_p_ corresponds to the maternal and paternal effects, respectively, if they are identifiable. In absence of family data, these effects cannot be distinguished and therefore cannot be estimated. To solve this problem, Hoggart et al. [20] proposed an appealing methodology to allow the detection of POE in genotype data of unrelated individuals only. By rewriting model (2) as:

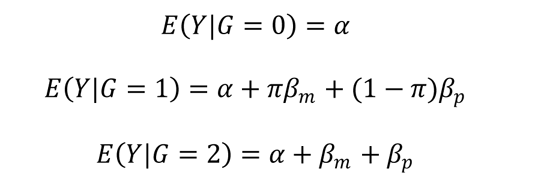

where π is a random variable following a Bernoulli distribution with parameter ½ (50% of alleles coming from the paternal and 50% from the maternal transmission), they observed that, in presence of POE, the phenotypic variance in heterozygous individuals should be higher than the phenotypic variances in the two groups of homozygotes:

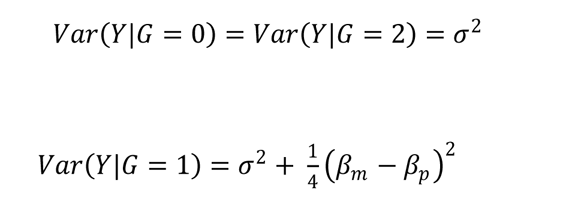

They then proposed to use the Brown-Forsythe (BF) test [31], a robust version of the Levene test, to test whether the phenotypic variance in heterozygote carriers of a given SNP is significantly higher than the phenotypic variances observed in homozygotes. They further showed that this BF test is equivalent to performing a linear model where the absolute deviation of the phenotype from the intra-genotype median is regressed on a binary variable indicating whether an individual *i* is heterozygote or not at the tested *j* SNP.

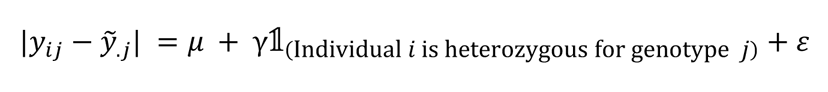

A positive and significant value for the γ regression coefficient associated to this indicator variable is a sign for POE.

They implemented this BF framework in the Quicktest program [https://wp.unil.ch/sgg/program/quicktest/] that can easily be applied to large GWAS genotype datasets to detect POE acting on a quantitative trait. Of note, as highlighted by Hoggart et al, while the presence of POE can lead to significant BF test, the inverse is not necessary true as a significant BF test can also be due to other phenomena such as haplotype effects, gene x gene or gene x environment interactions.

For the present work, all SNPs with imputation quality r^2^ > 0.5, minor allele frequency > 0.005, and a number of heterozygous individuals greater than 20, were tested through the BF methodology in relation to FV activity. Analyses were adjusted for age, sex, and genetically derived principal components. Additional adjustment on case-control status was performed in RETROVE.

The Quicktest software was applied in each study and results were then meta-analyzed using a fixed-effect meta-analysis as implemented In the GWAMA software [32].

Heterogeneity across study populations was assessed by the Cochran’s Q statistic and the I^2^ index.

Genome wide statistical significance was considered at BF p-values <5 10^-8^.

#### Search for gene x gene interactions

To further investigate the possible source explaining each genome–wide significant BF signal, we sought for SNPs that could modulate FV activity differentially according to the heterozygote status at the lead SNP identified by the QuickTest analysis. For this, in each contributing study population, we conducted a genome-wide interaction analysis based on a linear model where FV activity were regressed for age, gender, genetically derived principal components, the heterozygote status at the lead BF SNP, any SNP, and an interaction term between the latter two. All SNPs with imputation quality r^2^ < 0.5 and minor allele frequency < 0.005 were excluded from the analyses. These genome wide interaction analyses were conducted using Plink2 (www.cog-genomics.org/plink/2.0/) [33].

For each tested SNP, interaction terms were then meta-analyzed across the 4 studies using a fixed-effect meta-analysis as implemented in the GWAMA software [32].

## Results

In total, 4,505 individuals were studied in this work. A brief description of the general characteristics of the four contributing studies is given in Table 1.

**Table 1 :** Brief description of the studied populations.

|  | LURIC | MARTHA | MEGA | RETROVE |
| --- | --- | --- | --- | --- |
| N | 1,833 | 1,011 | 865 | 796 |
| % of VT cases | 0% | 100% | 100% | 50% |
| Age, years (SD) | 62.3 (10.8) | 47.6 (15.7) | 47.6 (12.9) | 54.4 (19.9) |
| Female sex, N (%) | 537 (29%) | 633 (63%) | 444 (51%) | 406 (51%) |
| Factor V (U/dL) | 1.13 (0.22) | 1.07 (0.23) | 0.95 (0.19) | 0.99 (0.20) |
| Platelets count | 231.1 (66.7) | 256.4 (68.9) | NA | 235.3 (62.4) |
| Neutrophil counts* | 59.6 (9.46) | 61.1 (8.91) | NA | 58.1 (9.7) |
| Smoking at sampling (%) | 1,163 (65%) | 167 (18%) | NA | 129 (16%) |

\*percentage compared to total white blood cells. In LURIC, smoking status includes both active and former smokers.

7,300,264 SNPs were tested in relation with FV activity through the BF framework. A Manhattan plot summarizing the statistical findings is shown in Figure 1. The associated Quantile-Quantile plot is given in Supplementary Figure S1. One locus, chr8q21.3, reached the prespecified genome wide statistical threshold of 5 x 10^-8^. The lead SNP was rs75463553 and its POE γ coefficient was 0.128±0.022 (p = 3.45 x 10^-9^). As shown in Table 2, the POE γ coefficients were very homogeneous across the 4 contributing studies as were the allele frequencies. We observed that the variance in FV activity was higher in carriers of the G/T genotype compared to the combined groups of G/G and T/T genotypes while no association with mean FV levels was observed (p=0.49 in the combined 4 studies).

**Table 2 :** Association of rs75463553 with phenotypic mean and variance of FV activity.

|  | LURIC | MARTHA | MEGA | RETROVE |
| --- | --- | --- | --- | --- |
| N | 1,833 | 1,011 | 865 | 796 |
| Minor Allele frequency (G/T) | 0.145 | 0.117 | 0.156 | 0.106 |
| Imputation $r^2$ | 0.980 | 0.960 | 0.975 | 0.982 |
| N |  |  |  |  |
| GG | 1338 | 779 | 614 | 635 |
| GT | 463 | 222 | 235 | 153 |
| TT | 32 | 10 | 16 | 8 |
| FV activity means |  |  |  |  |
| GG | 1.132 | 1.056 | 0.958 | 0.990 |
| GT | 1.142 | 1.097 | 0.946 | 0.998 |
| TT | 1.059 | 1.082 | 0.912 | 1.119 |
| FV activity standard deviation |  |  |  |  |
| GG | 0.212 | 0.224 | 0.176 | 0.192 |
| GT | 0.228 | 0.264 | 0.217 | 0.220 |
| TT | 0.185 | 0.261 | 0.157 | 0.292 |
| POE $\gamma$ effect | 0.098±0.033 | 0.181±0.049 | 0.152±0.050 | 0.115±0.055 |
| P | $1.693 \times 10^{-3}$ | $1.042 \times 10^{-4}$ | $1.307 \times 10^{-3}$ | $1.913 \times 10^{-2}$ |

EAF: Estimated allele frequency
The POE $\gamma$ effects did not show any evidence for heterogeneity across cohorts ( $I^2=0$ , $P = 0.518$ )

**Figure 1 :**
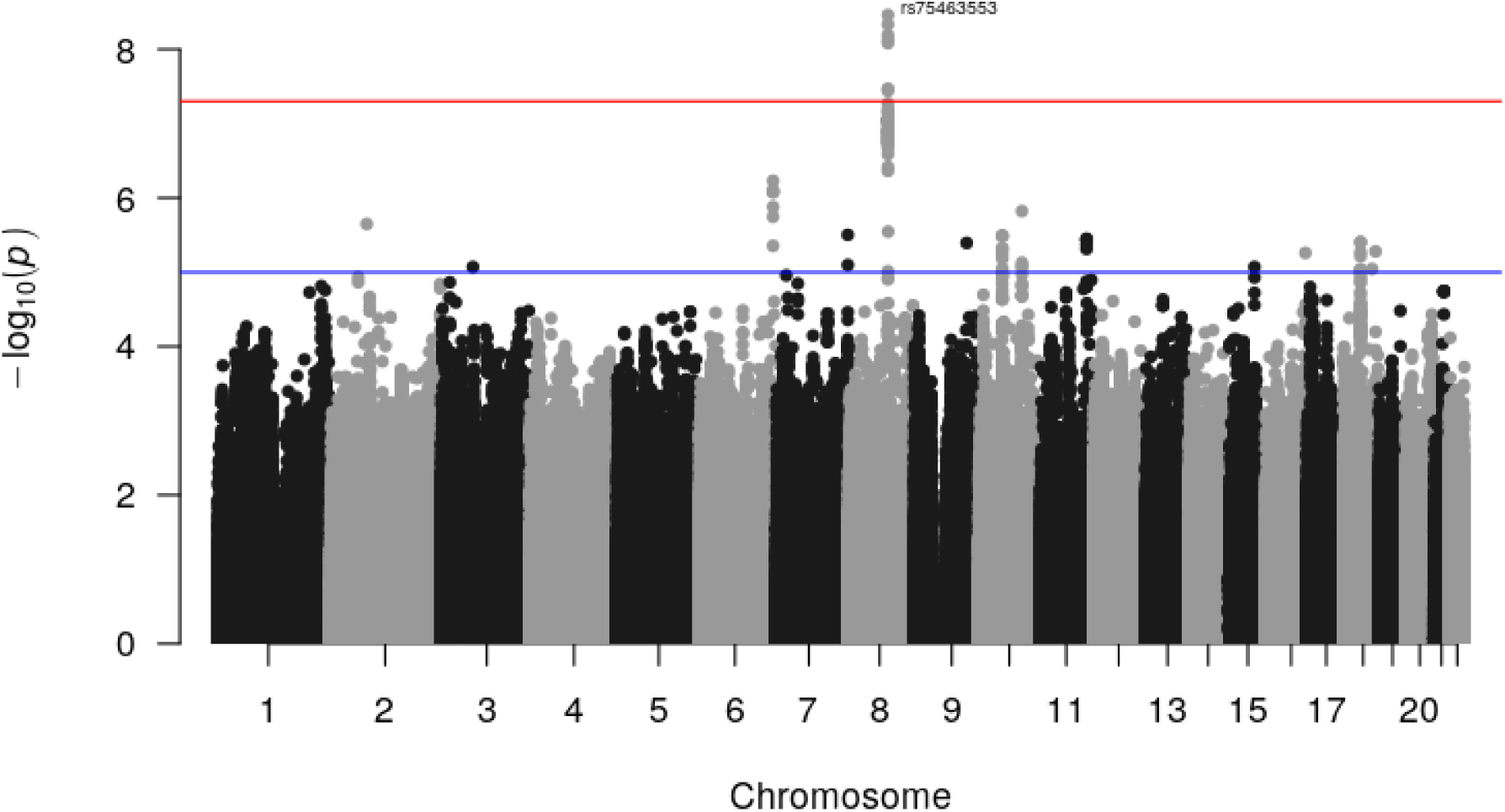
Manhattan plot of 4 studies of unrelated Europeans individuals for detecting POE in FV plasma level (n=4,505).

rs75463553 maps to an intronic region of the non-coding RNA LOC105375623, downstream to *SLC7A13* and upstream to *ATP6V0D2*. It is in strong linkage disequilibrium (LD) with 6 other nearby SNPs with genome wide significant BF p-value (Supplementary Figure S2 – Supplementary Table S2). These 6 SNPs generate 3 haplotypes with frequency > 1%. None of them associated with mean FV activity (Supplementary Table S3) suggesting that the detected BF signal was unlikely due to LD effects between nearby SNPs.

By interrogating various online genomic resources, we observed that:

- rs75463553 is an expression quantitative trait locus (eQTL) for LOC105375623 in testis but also for nearby *SLC7A13* (testis) and *WWP1* (cultured fibroblasts) genes (https://gtexportal.org/home/snp/rs75463553). It is also moderately associated with the expression of more distant genes on 8q21.3 such as *RMDN1*, *CPNE3* and *NTAN1P2*.

- rs75463553 also associated (p = 1.76 x 10^-8^) with plasma levels of RMDN1 protein in the Fenland study [34], but it was mainly a protein quantitative trait locus (pQTL) (p

= 2.92 x 10^-13^) for Carbonic anhydrase XIII whose structural gene (*CA13*), located on 8q21.2, is more than 1 Mb from rs75463553.

- While moderately associated (p = 5.9 x 10^-7^) with whole blood DNA methylation levels at *CA13* cg0571334 CpG site in the GoDMC database (http://mqtldb.godmc.org.uk/index) [35], rs75463553 demonstrated strong associations with whole blood DNA methylation at several CpG sites at the *PSKH2* locus (Supplementary Figure S3) such as cg00001099 (p = 5.4 x 10^-291^), cg26186954 (p = 7.54 x 10^-185^) and cg20982735 (p = 1.13 x 10^-157^).

*PSKH2* is located upstream *ATP6V0D2* gene and has been shown to harbor SNPs associated with neutrophil count [36] whose expression has been shown to correlate with that of *F5* in individuals with inflammatory disorders [37]. We then investigated the association of rs75463553 with neutrophil counts in LURIC, MARTHA and RETROVE. We observed a trend for neutrophil counts being higher in homozygote carriers of the rs75463553-T allele (Table 3). However, this association was mainly observed in individuals with high platelets counts. As shown in Table 3, when the combined sample was divided according to the median of platelets observed in the global population, rs75463553-TT carriers with platelets count over the median exhibited significantly (p = 0.0023) higher neutrophil count while no association (p = 0.59) was observed in the group of individuals with lower platelets. Surprisingly, we further observed that the significant association was mainly restricted to smokers (Table 4). However, no such interactive effects were observed on FV activity (data not shown) suggesting that the complex relationship between rs75463553, platelets and neutrophils count would unlikely explain the statistical BF signal observed on FV activity. Consistent with this hypothesis is the observation that the effect of rs75463553 on FV variability, as assessed by the BF methodology, still holds according to platelets and smoking (Supplementary Table S4), except in smokers with high platelet counts.

**Table 3 :** Association of rs75463553 with neutrophil counts in LURIC, MARTHA and RETROVE studies according platelets count.

|  | All population |  |  | Platelets<=230 |  |  | Platelets>230 |  |  |
| --- | --- | --- | --- | --- | --- | --- | --- | --- | --- |
|  | LURIC | MARTHA | RETROVE | LURIC | MARTHA | RETROVE | LURIC | MARTHA | RETROVE |
| GG | 4.21 (1.54)<br>N = 1312 | 3.95 (1.43)<br>N = 675 | 3.93 (1.54)<br>N = 635 | 3.89 (1.38)<br>N = 728 | 3.62 (1.24)<br>N = 239 | 3.67 (1.39) N = 318 | 4.60 (1.63)<br>N = 584 | 4.13 (1.49)<br>N = 436 | 4.19 (1.64)<br>N = 317 |
| GT | 4.17 (1.57)<br>N = 455 | 4.08 (1.59)<br>N = 186 | 3.93 (1.64)<br>N = 153 | 3.92 (1.52)<br>N = 255 | 3.59 (1.52)<br>N = 72 | 3.54 (1.33)<br>N = 73 | 4.49 (1.58)<br>N = 200 | 4.38 (1.56)<br>N = 114 | 4.29 (1.82)<br>N = 80 |
| TT | 4.43 (2.11)<br>N = 33 | 4.87 (2.79)<br>N = 10 | 3.75 (1.51)<br>N = 8 | 3.71 (0.94)<br>N = 21 | 4.02 (0.94)<br>N = 3 | 3.19 (0.37)<br>N = 6 | 5.68 (2.94)<br>N = 12 | 5.24 (3.30)<br>N = 7 | 5.43 (2.79)<br>N = 2 |
| $\beta \pm SE$ | $\beta = 0.26 \pm 0.27$ | $\beta = 0.92 \pm 0.47$ | $\beta = -0.16 \pm 0.35$ | $\beta = -0.18 \pm 0.32$ | $\beta = 0.42 \pm 0.76$ | $\beta = -0.24 \pm 0.35$ | $\beta = 1.09 \pm 0.48$ | $\beta = 1.09 \pm 0.59$ | $\beta = 0.53 \pm 0.76$ |
| P* | P = 0.351 | P = 0.053 | P = 0.654 | P = 0.562 | P = 0.581 | P = 0.497 | p = 0.023 | P = 0.063 | P = 0.480 |
| Combined $\beta$ | | $\beta = + 0.31 \pm 0.22$<br>P = 0.146 | | | $\beta = -0.137 \pm 0.25$<br>P = 0.59 | | | $\beta = + 1.08 \pm 0.35$<br>P = 0.0023 | |

\* Association was tested using a linear model adjusted for age and sex under the assumption of recessive effect $\beta$

**Table 4 :** Association of rs75463553 with neutrophil counts according to platelets counts and smoking in LURIC, MARTHA and RETROVE studies combined.

|  | Platelets≤230 |  | Platelets>230 |  |
| --- | --- | --- | --- | --- |
|  | Non Smokers | Smokers | Non Smokers | Smokers |
| rs75463553 |  |  |  |  |
| GG/GT | 3.59 (1.30)<br>N = 945 | 4.04 (1.47)<br>N = 741 | 4.1 (1.52)<br>N = 1044 | 4.77 (1.65)<br>N = 688 |
| TT | 3.49 (0.83)<br>N = 16 | 3.81 (0.92)<br>N = 14 | 4.19 (1.5)<br>N = 11 | 6.97 (3.44)<br>N = 10 |
| $\beta \pm SE$ | $\beta = -0.032 \pm 0.323$ | $\beta = -0.227 \pm 0.394$ | $\beta = +0.084 \pm 0.457$ | $\beta = +2.17 \pm 0.538$ |
| P* | P = 0.921 | P = 0.564 | P = 0.853 | P = $6.07 \cdot 10^{-5}$ |

\* Association was tested using a linear model adjusted for age, sex and cohort under the assumption of recessive TT effect

Of note, in LURIC, MARTHA and RETROVE where neutrophils and platelets count were measured, FV activity did not exhibit significant correlation with them (Supplementary Table S5).

We then further explored whether the original detected BF signal could be explained by the interaction of rs75463553 with other SNPs. The genome wide scan conducted in the four contributing studies identified one genome wide significant (p = 2.6 10^-8^) interaction (Supplementary Table S6, Supplementary Figure S4). In heterozygous carriers of the rs75463553-T allele, carrying the rs7190785-A allele at *GRIN2A* on chromosome 16p13.2 was associated with increased FV activity (β = 0.05 ± 0.01, p = 2.52 10^-8^). By contrast, no association was observed for the rs7190785 allele in individuals with GG or TT genotypes at rs75463553.(β = -0.01 ± 0.01, p = 0.22). This phenomenon was consistent in LURIC, MARTHA and MEGA, but not in RETROVE (Supplementary Table S7). It is worthy to note that a second interaction signal nearly reached genome wide significance (p = 6.27 x 10^-8^) mapping to *POM121L12* locus on chr7p12.1 which, as *GRIN2A*, has been observed in several GWAS to associate with smoking phenotypes [38–40].

## Discussion

This work was motivated by the search of non-additive genetic effects that could contribute to the inter individual variability of FV plasma activity. To achieve this objective, we used an underestimated methodology, but with great potential, that allows to leverage on existing GWAS data in a very efficient and quick way as it is implemented in the easy-to-use Quicktest software [20]. Even if the method was initially proposed to detect POE effects, it also has potential to detect non-additive genetic effects that could be due to gene x gene or gene x environment interactions. Its application here provides evidence for the presence of gene x smoking interaction in the modulation of FV plasma activity.

The application of this methodology on 4,505 individuals phenotyped for FV activity and with GWAS data identified SNPs at the 8q21.3 locus significantly associated with the variability of FV activity. Using publicly available resources, we observed that the lead SNP at this locus, rs75463553, associates with DNA methylation levels of this locus, in particular with several CpG sites at the *PSKH2* gene. *PSKH2* encodes a protein serine kinase about which, little is known. Some genetic studies have linked *PSKH2* SNPs with neutrophil [36,41] and myeloid leukocyte [36] counts whose role in the coagulation and thrombotic pathways have been highly discussed in the literature [42]. In our work, we did observe an association between rs75463553 and neutrophil counts, but this association could not explain the genome-wide signal we detected. We sought to investigate whether this signal could be caused by POE in a family study but we were only able to assess this hypothesis in a sample of 21 families from the GAIT1 study [43]. Unfortunately, only 26 informative meioses were available to test for a differential paternal – maternal effect of the rs75463553 and no statistical association was observed (p>0.50) (data not shown). Of note, *PSKH2* was not detected to be prone to POE using an alternative methodology based on sequencing data and applied to several phenotypes, including neutrophil counts, from the UKbiobank [44]. This would suggest that the signal we detected using the QuickTest software could be due to other phenomena rather than POE. In line with this hypothesis are the candidate gene x gene interactions we identified with two loci,

*GRIN2A* and *POM121L12*, that have been proposed to be involved in smoking phenotypes in previous GWAS [38–40]. Unfortunately, smoking status at the time of blood sampling was not available in all contributing studies and it was then not possible to assess whether the observed signal was due to complex interactions between several polymorphisms and smoking. Similarly, our studies had very limited information about additional environmental covariates that prevented us from performing more exhaustive gene x environment interaction analyses and from determining whether the *PSKH2* locus statistical signal could underline such epistasis phenomena.

In conclusion, this work provides strong statistical argument supporting the role of the *PSKH2* locus in the variability of FV activity. But more in-depth investigations are now needed to characterize the exact underlying mechanisms. Besides, this work also emphasizes how to leverage on existing large GWAS datasets to detect non additive allele effects using the Brown-Forsyth methodology as implemented in the QuickTest program which could explain part of the missing heritability still existing for most complex phenotypes.

## Supplementary Material

Tables S1-S7

Figures S1-S4

