## Supplementary Figures for "A genome wide search for non-additive allele effects identifies *PSKH2* as involved in the variability of Factor V activity"

**Supplementary Material**

**Figure S1**

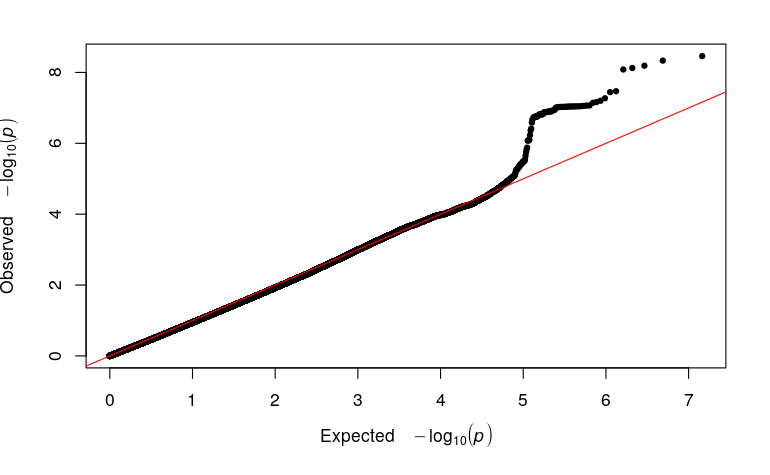

Supplementary Figure S1 : Quantile-Quantile plot representation of the meta-analysis results

**Figure S2**

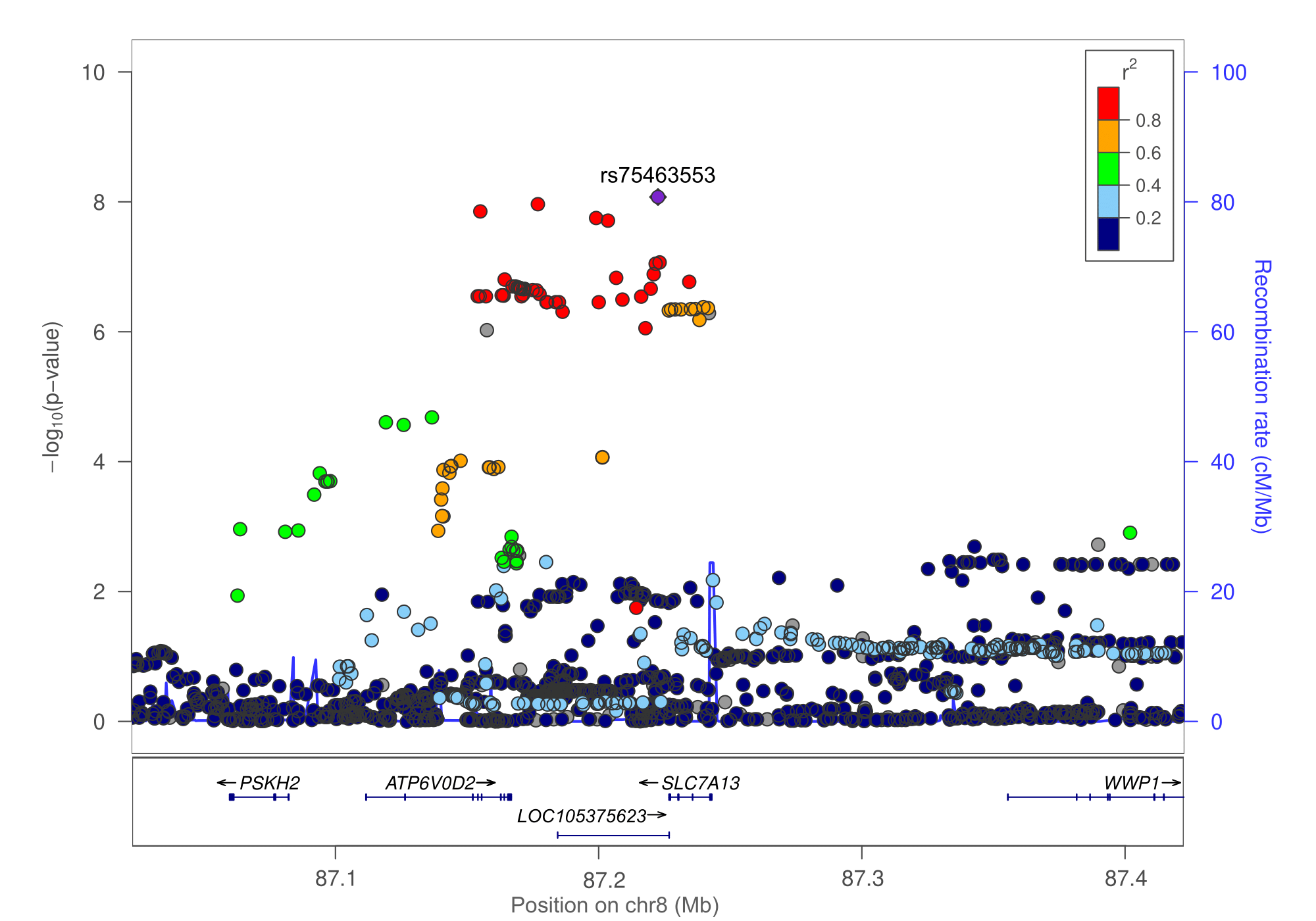

Supplementary Figure S2 : Regional plot describing the Brown-Forsythe p-values observed at the *SLC7A13* locus

**Figure S3**

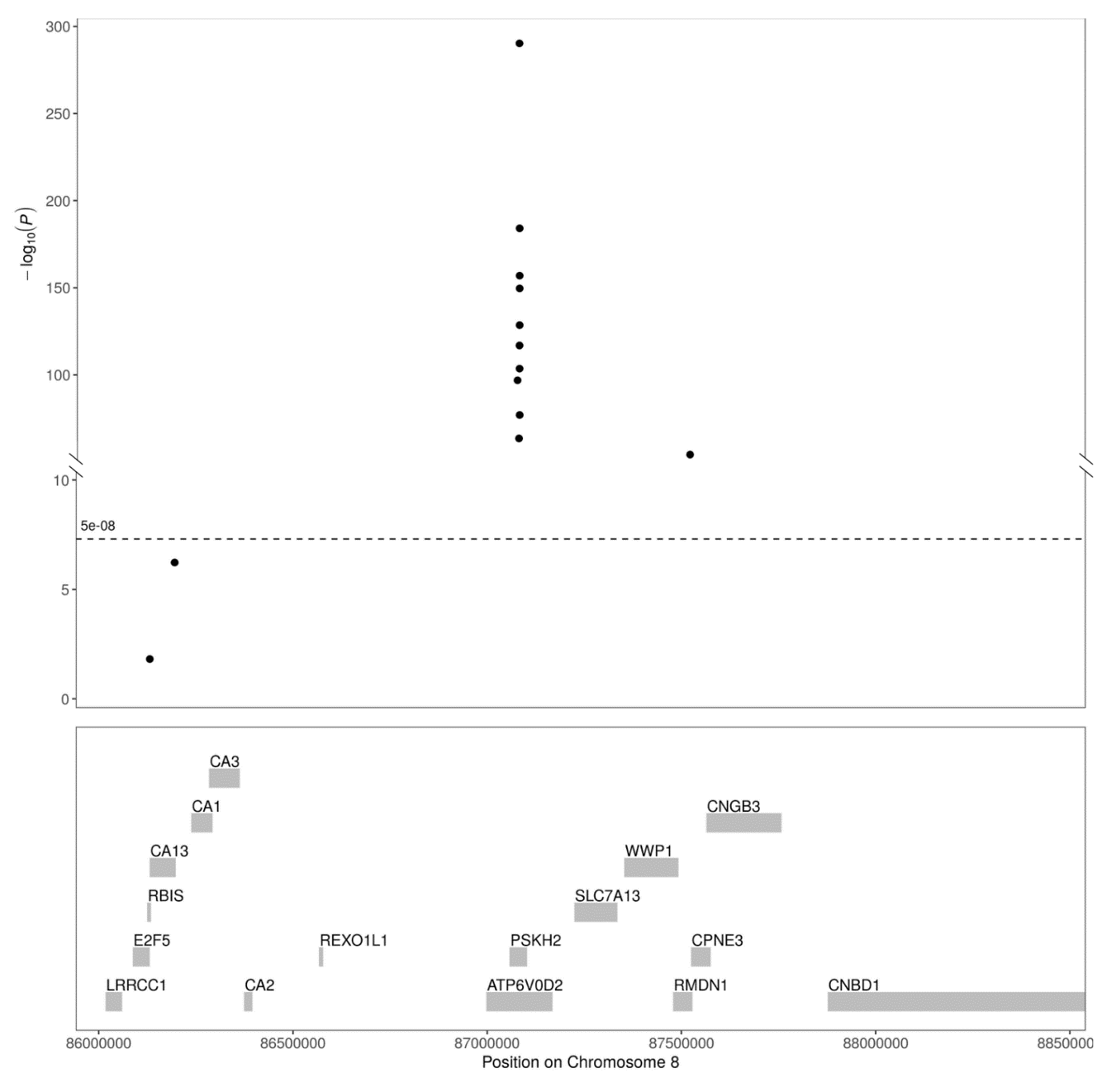

Supplementary Figure S3 : Association of rs75463553 with CpG sites at the *PSKH2* locus derived from GoDMC portal (mqtldb.godmc.org.uk/index)

**Figure S4**

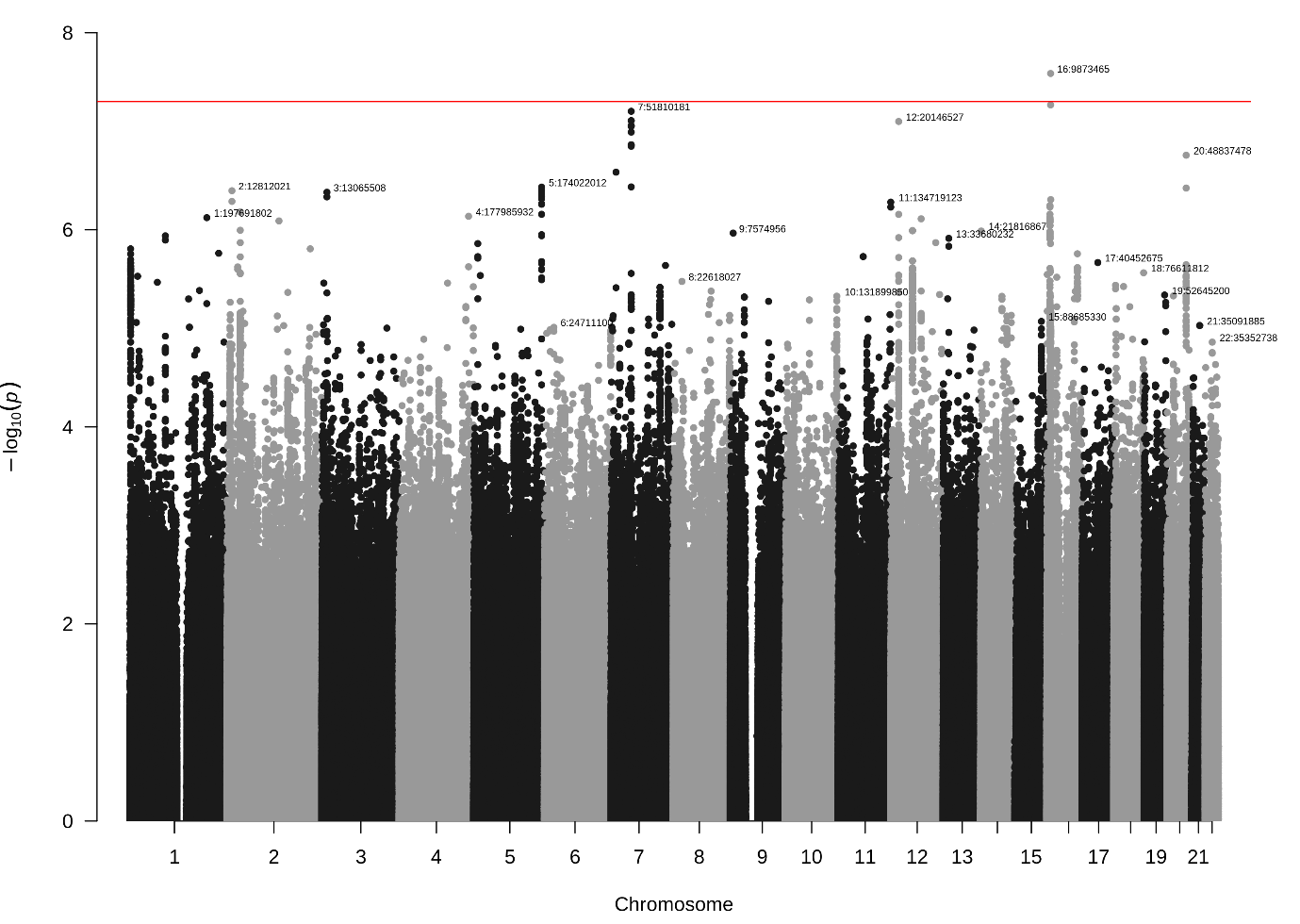

Supplementary Figure S4 : Manhattan plot representation of the genome-wide search for rs75463553 x SNP interactions on FV activity resulting for the meta-analysis of 4 independent studies
